# Improving MALDI Mass Spectrometry Imaging Performance: Low-Temperature Thermal Evaporation for Controlled Matrix Deposition and Improved Image Quality

**DOI:** 10.1101/2025.01.06.631532

**Authors:** Toufik Mahamdi, Cristina Gomez Serna, Roger Giné, Jordi Rofes, Shad Arif Mohammed, Pere Ràfols, Xavier Correig, María García-Altares, Carsten Hopf, Stefania-Alexandra Iakab, Oscar Yanes

## Abstract

The deposition of matrix compounds significantly influences the effectiveness of matrix-assisted laser desorption/ionization (MALDI) Mass Spectrometry Imaging (MSI) experiments, impacting sensitivity, spatial resolution, and reproducibility. Dry deposition methods offer advantages by producing homogeneous matrix layers and minimizing analyte delocalization without the use of solvents. However, refining these techniques to precisely control matrix thickness, minimize heating temperatures, and ensure high-purity matrix layers is crucial for optimizing MALDI-MSI performance. Here, we present a novel approach utilizing low-temperature thermal evaporation (LTE) for organic matrix deposition under reduced vacuum pressure. Our method allows for reproducible control of matrix layer thickness, as demonstrated by linear calibration for two organic matrices, 2,5-dihydroxybenzoic acid (DHB), and 1,5-Diaminonaphthalene (DAN). The environmental scanning electron microscopy images reveal a uniform distribution of small-sized matrix crystals, consistently on the submicrometer scale, across tissue slides following LTE deposition. Remarkably, LTE serves as an additional purification step for organic matrices, producing very pure layers irrespective of initial matrix purity. Furthermore, stability assessment of MALDI-MSI data from mouse brain sections coated with LTE-deposited DHB or DAN matrix indicates minimal impact on ionization efficiency, signal intensity, and image quality even after storage at - 80°C for two weeks, underscoring the robustness of LTE-deposited matrices for MSI applications. Comparative analysis with the spray-coating method highlights several advantages of LTE deposition, including enhanced ionization, reduced analyte diffusion, and improved MSI image quality.

## 1. Introduction

Since its introduction by Caprioli et al. in the late 1990s^1^, mass spectrometry imaging (MSI) has become a powerful tool for analyzing the spatial distribution of various molecular classes within biological tissues. Matrix-Assisted Laser Desorption/Ionization-Mass Spectrometry Imaging (MALDI-MSI) enables the visualization of compounds including proteins^2^, lipids^3^, metabolites and drugs^4^ within tissue sections. Matrix deposition stands out as a crucial step in sample preparation, significantly impacting the quality of ion images produced.

Various techniques are employed for applying matrices in tissue samples, including wet methods like spraying^5,6^ and dry methods like sublimation^7,8^. Sublimation offers advantages over spraying in terms of matrix deposition homogeneity^7,9,10^, analyte delocalization^11,12^, and spatial resolution^7-9,13^. Devices like the HTX TM-Sprayer and SunCollect MALDI-Sprayer are used for wet deposition, while iMLayer and HTX sublimator are utilized for dry deposition. However, current devices do not adequately address challenges like controlling matrix thickness, preventing biological degradation from elevated temperatures, and ensuring the purity of matrix deposition.

In this regard, thin film deposition techniques like physical vapor deposition (PVD) and chemical vapor deposition (CVD) have been refined for precise deposition of pure and thin material layers. PVD encompasses processes like vacuum or thermal evaporation (TE), sputtering, and ion plating, which involve vaporizing atoms or molecules from a source, transporting them through a vacuum, and condensing them onto a substrate. PVD is widely used in various industries to enhance properties such as hardness and wear resistance^14-18^. In MALDI-MSI, PVD coating with gold or silver nanoparticles has been explored as an alternative to organic matrices^19,20^. TE techniques offer advantages like higher purity films and better control over film thickness^21^. Researchers have investigated factors like substrate holder geometry and vacuum pressure to enhance coating uniformity and distribution. Low temperature evaporation (LTE) has emerged as a notable advancement in vacuum evaporation techniques, particularly beneficial for organic materials sensitive to temperature variations. LTE finds applications in diverse industries, including nanotechnology, solar cells, and pharmaceuticals^22-24^.

In this study, we propose using LTE as a novel dry deposition method for applying organic matrix compounds in MALDI-MSI to achieve better control over the matrix layer and ensure a high-quality, uniform matrix crystal coating. The evaporator system, equipped with LTE sources, facilitates precise deposition at low evaporation temperatures and high-vacuum conditions (<5×10^−5^ mbar). Our reproducible LTE deposition method for DHB and DAN matrices at 80°C ensures controlled deposition rates while minimizing sample degradation. ESEM analysis confirms the homogeneous distribution of submicrometer-sized matrix crystals over tissue samples. Notably, using <99% pure matrices does not compromise lipid analyte ionization or MS image quality compared to >99% pure matrices. Tissue sections coated with DAN or DHB matrix stored at -80°C for 2 weeks yield consistent MALDI-MSI results. Comparisons with the traditional spray-coating method demonstrate LTE’s superiority in detecting more lipids, yielding higher signal intensities, and producing higher-quality MALDI images with reduced analyte diffusion.

## 2. Materials and methods

### 2.1. Reagent and chemicals

Ethanol (HPLC grade) was purchased from Scharlab S.L. 2,5-Dihydroxybenzoic acid (DHB) ≥99.0%, DHB 98%; DAN: 1,5-Diaminonaphthalene (DAN) ≥99.0%, and DAN 97% were purchased from Sigma-Aldrich. Microscope slides were purchased from Epredia and Conductive ITO-coated slides were purchased from Bruker.

### 2.2. Tissue Sampling

Bovine liver procured from a local butcher shop was stored in a -80°C freezer and subsequently transferred to a +4°C chamber for preparation. The liver was dissected into small pieces and placed in micro-tubes containing three metal beads for homogenization. Homogenization of the liver tissue was achieved using the FastPrep-24 homogenizer (M.P. Biomedicals, Irvine, California, USA) with the speed set to 4 rpm for 15 seconds. Following homogenization, the metal beads were removed, and the micro-tubes were immediately immersed in ice-cold water and subsequently transferred to a -80°C freezer for storage. Brain tissues utilized in this study were harvested from C57BL/6 mice, promptly flash-frozen on dry ice, and stored at -80°C until further processing. Fresh-frozen mouse brain and liver homogenate samples were sectioned at -20°C into 10 µm thick sections using a CryoStar NX50 cryostat. Subsequently, the tissue sections were mounted onto indium tin oxide (ITO-coated) microscope slides and promptly stored in a -80°C freezer for subsequent analysis.

### 2.3. Development of a matrix deposition method by low thermal evaporation and thickness layer characterization

The thermal evaporator nanoPVD-T15A (Moorfield Nanotechnology, United Kingdom) utilized in this study is equipped with two low-temperature evaporation (LTE) sources featuring shutters and a quartz crystal sensor head (Supplementary Figure S1). These LTE cells contain crucibles enabling the deposition of volatile materials at temperatures up to 600°C. All components, including temperature, power supply, shutters, and substrate holder rotation, are conveniently controlled via a touchscreen Human-Machine Interface (HMI). The chamber is integrated with a turbomolecular pumping system, and access is facilitated through a hinged lid, revealing a stage suitable for accommodating substrates up to 10.16 cm (4”) in diameter.

The thermal evaporator features a quartz crystal microbalance sensor for monitoring thickness, relying on parameters such as density and acoustic number of the organic matrices. Density and acoustic number values were set at 1.4 and 0.1, respectively, for both DHB and DAN matrices, as reference data for these parameters were unavailable. The optimal evaporation temperature for DHB and DAN was determined experimentally under a working vacuum pressure of 5×10^−5^ mbar. This temperature corresponds to the point at which the deposition rate peaks and then stabilizes, as shown in Supplementary Figure S2. Deposition of DHB and DAN matrices involved placing 300 mg of the respective matrix into the crucible within the LTE source. Upon chamber evacuation, the LTE source was heated incrementally from 55°C to 80°C, and the substrate shutters were opened after reaching 80°C to enable deposition of evaporated matrix particles onto the sample slide. Deposition ceased by closing all the shutters upon reaching the targeted matrix layer thickness.

Calibration of DHB and DAN matrix thicknesses against the thickness monitored by the thermal evaporator sensor was conducted using silicon wafer slides for real thickness measurements and glass slides for weight measurements. Calibration points were selected at intervals of 2500 Å for DHB and 2000 Å for DAN, spanning from 5000 Å to 17500 Å for DHB and from 4000 Å to 12000 Å for DAN. Silicon wafers were prepared by cutting into slides of 5×1 cm dimensions, mounted on glass slides, and subsequently placed on the substrate holder of the thermal evaporator. Up to three slides (75×25 cm) could be accommodated on the substrate target, allowing for two replicates of matrix layer deposition on silicon slides and one glass slide sample per experiment (Supplementary Figure S3). Each calibration point was replicated six times, with three experimental depositions performed on the same day.

Electron microscopy measurements, specifically Environmental Scanning Electron Microscopy (ESEM) analysis (FEI Quanta 600, FELMI-ZFE), were employed to determine the real thickness of the matrix layer deposited on silicon. Following matrix application, the silicon cross sections were observed using ESEM under low vacuum (0.68 Torr) utilizing a Large Field Detector (LFD). To determine the optimal matrix thickness for MALDI imaging of lipids, liver homogenate tissue sections were coated using thermal evaporation with varying DHB and DAN layer thicknesses. Six DHB layer thicknesses ranging from 5000 Å to 17500 Å and five DAN thicknesses from 4000 Å to 12000 Å were assessed.

### 2.3 Matrix purity analysis

For the analysis of the matrix alone, microscope slides were initially washed with ethanol and allowed to dry. Pure matrices (≥99% purity) of DHB and DAN, as well as non-pure matrices (98% DHB and 97% DAN), were deposited onto the microscope slides using two distinct approaches. The first method involved employing the developed thermal evaporation technique with controlled source and substrate shutters (referred to as “C_Sh”), while the second deposition was conducted with both shutters left open throughout the evaporation process (“O_Sh”).

For comparison experiments between pure matrices and non-pure matrices, sagittal sections (10 µm) of mouse brain tissues were utilized. The pure and non-pure matrices were deposited onto the brain sections using the optimized evaporation protocol.

### 2.4 Sample stability analysis

DAN and DHB matrices were deposited onto mouse brain sections using LTE following the optimized protocol outlined in Supplementary Methods. The coated brain slides were subsequently stored in a -80°C freezer. Sample slides were retrieved for analysis after storage durations of 1 hour, 24 hours, and 14 days prior to conducting MALDI-MSI and ESEM analysis.

### 2.5. Comparison between LTE and spray-coating

For the LTE samples, sagittal sections of mouse brains were coated with DAN and DHB matrices, subsequently packaged in dry ice and shipped to Germany. Upon arrival, they were promptly stored at -80ºC prior to MALDI-MSI analysis.

For the spray-coating samples, DHB (40 mg/mL in 70% methanol) and DAN (10 mg/mL in 50% acetonitrile) were sprayed onto the samples using an HTX-M5 sprayer (HTX Technologies, LLC, Chapel Hill, USA). The spray parameters were as follows: For DHB, the temperature was set to 65°C, with 11 passes and a flow rate of 0.05 µL/min; for DAN, the temperature was set to 60°C, with 10 passes and a flow rate of 0.06 µL/min.

### 2.4. MALDI imaging spectra acquisition

#### Two MALDI instruments were employed in this study

A dual-ion funnel MALDI/ESI injector (Spectroglyph, Kennewick, WA) equipped with an Orbitrap Exploris 120 mass spectrometer (Thermo Fisher Scientific) was utilized for MALDI MS imaging analysis, with a spatial step-size of 20 and 30 μm. Spectra were acquired in the m/z mass range of 400 to 1200 Da for MALDI imaging of lipids and from 50 to 600 Da for the analysis of only matrix samples at a mass resolution of 60,000 (FWHM at m/z 200). Processing of MSI datasets, as well as MSI image reconstruction and visualization, were carried out using the rMSI2 R package (https://github.com/prafols/rMSI2). The identified MALDI ions were cross-referenced with the Lipid Maps database (https://www.lipidmaps.org/). A mass tolerance of +/-0.005 m/z was applied for the search, and [M+H]^+^, [M-H2O+H]^+^, [M+Na]^+^, and [M+K]^+^ adducts were selected for positively detected ions, while [M-H]^-^ adducts were selected for negative ions.

A timsTOFfleX mass spectrometer (Bruker Daltonik, Bremen, Germany), operated in tims OFF mode, was also utilized. The mass range was set between m/z 400 and 1200, with a spatial step-size of 10 and 30 μm. Between 50 and 200 laser shots were summed up per pixel for all experiments, with the laser operating at a repetition rate of 10 kHz, except for some exceptions at 5 kHz. All raw data were directly uploaded and processed in SCiLS lab (Version 2023a Pro). The displayed data, including ion images and spectra, were normalized to the root mean square (RMS).

## 3. Results and discussion

### 3.1. Control and reproducibility of matrix layer thickness by low-temperature thermal evaporation

Given the lack of prior research on LTE-based organic matrix deposition for MALDI-MSI, our initial aim was to determine the optimal evaporation temperature for matrices DHB and DAN, which we found to be 80°C under a vacuum pressure of 5×10^−5^ mbar. Unlike sublimation methods requiring higher temperatures (120-140°C for DHB^7,8^ and 110-140°C for DAN^8,25^) and additional cooling of sample slides using ice or a dry ice bath to ensure homogenous deposition, the LTE method operates at lower evaporation temperatures. By opening the shutters only after reaching this optimal temperature, the LTE approach ensures that only evaporated matrix particles reach the sample substrate, thereby minimizing the introduction of volatile impurities.

The thickness of the matrix layer is a critical parameter in MALDI imaging experiments as it determines the migration efficiency of analyte molecules through the tissue to the matrix layer surface. Hence, our primary aim was to develop a reproducible deposition method capable of controlling the deposited matrix thickness. The quartz sensor integrated into the thermal evaporator nanoPVD-T15A relies on the density and acoustic number of the deposited material to calculate thickness in angstroms (Å). Following the protocol outlined in Supplementary Methods, we deposited six layer thicknesses of DHB ranging from 5,000 to 17,500 Å and five layer thicknesses of DAN ranging from 4,000 to 12,000 Å.

Environmental Scanning Electron Microscopy (ESEM) images depicted in Figure 1a illustrate the characteristics of six different DHB layers. Notably, the deposition of DHB resulted in homogeneous layers with a sparse distribution of small crystals at the surface up to a monitored thickness of 10,000 Å. However, beyond this threshold, the number of matrix crystals increased proportionally with the monitored layer thickness, becoming more pronounced from 12,500 Å onwards. Conversely, the application of DAN matrix using LTE deposition yielded very homogeneous layers with smaller crystals at the surface (Figure 1d) compared to DHB. Furthermore, observations from ESEM images indicated that the matrix layer thickness reached a point of saturation at approximately 12,500 Å for DHB and 10,000 Å for DAN. Beyond these thresholds, additional deposited particles led to the formation of more crystals on the surface rather than contributing to the matrix layer.

**Figure 1:**
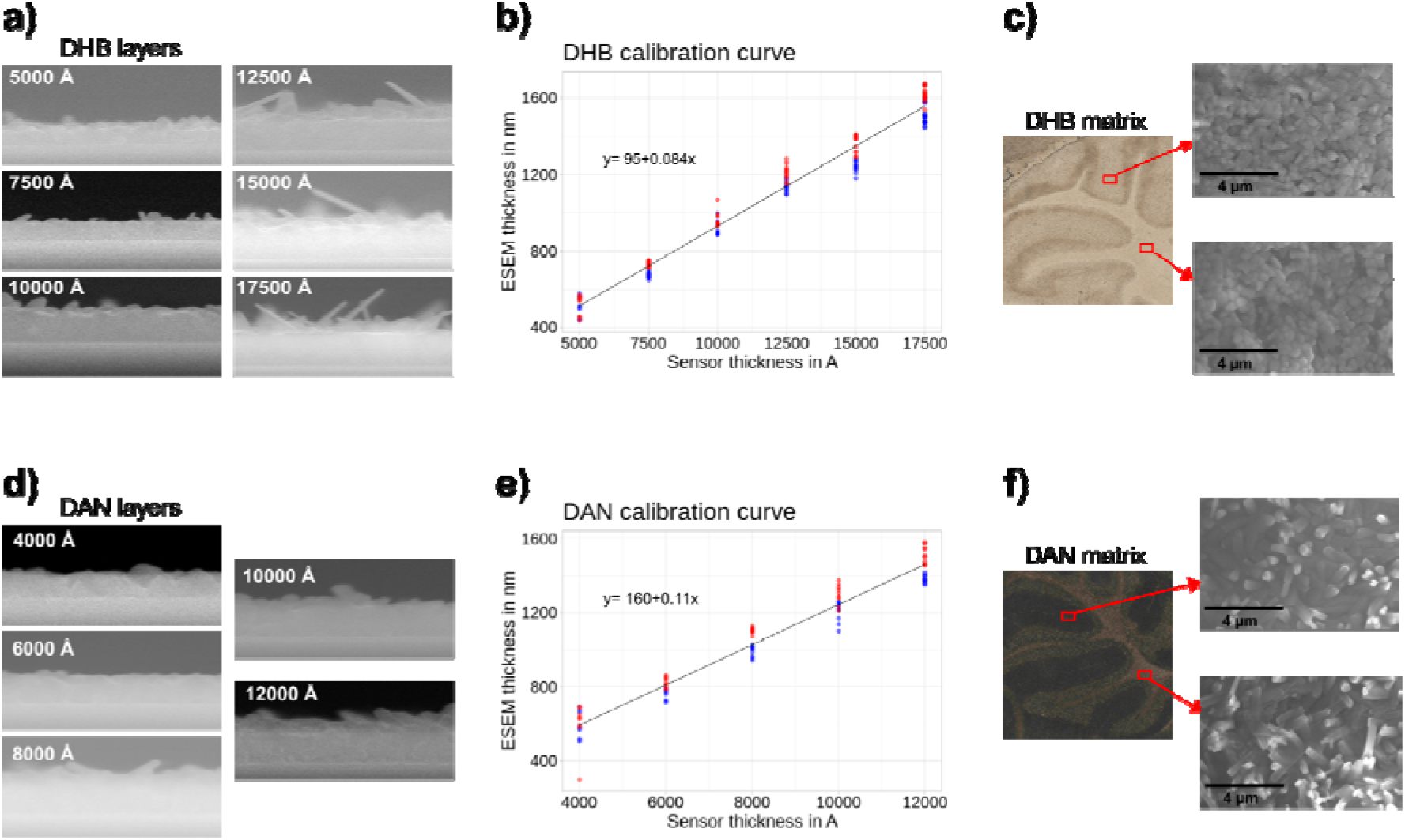
a) cross-section ESEM images of six layers of DHB (5,000 to 17,500 Å monitored by the quartz sensor). b) Calibration curve depicting the deposition of DHB, showing the relationship between the sensor thickness reported in angstrom units and the real thickness measured by ESEM in nanometers. c) Microscopy image of a mouse cerebellum tissue coated with DHB matrix (left) with corresponding ESEM images of a DHB surface layer acquired at 10,000 magnifications from the gray matter region (top right) and the white matter region (bottom right). d) cross-section ESEM images of five layers of DAN (4,000 to 12,000 Å monitored by the quartz sensor). e) Calibration curve illustrating the deposition of DAN, indicating the correlation between the sensor thickness reported in angstrom units and the real thickness measured by ESEM in nanometers. Red and blue points correspond to thickness measurements taken from the silicon slide positioned in the middle and top position of the substrate holder, respectively. f) Microscopy image of a mouse cerebellum tissue coated with DAN matrix (left) with related ESEM images of a DAN surface layer acquired at 10,000 magnifications from the gray matter region (top right) and the white matter region (bottom right). Scale bar, 4 µm.

Moreover, three different targeted thicknesses were deposited on the same day, and six replicates were applied on different days for each calibration point. Results demonstrated that the LTE protocol described herein can produce highly repetitive and reproducible matrix layers with low standard deviation. Additionally, we calculated the calibration curves of DHB (y=95+0.084x) and DAN (y=160+0.11x) matrices, where the real thickness measured by ESEM (y) exhibited a linear relationship with the sensor-monitored thickness (x) for both matrices, showing strong correlation coefficients (R^2^= 0.99) (Figure 1b and 1e). We also derived calibration curves for the deposited matrix weight (µg/cm^2^), resulting in equations y= 13+0.013x for DHB and y= 6.7+0.013x for DAN (Supplementary Figures S4 and S5).

### 3.2. Investigation of matrix crystal formation

We investigated the matrix crystals formed by LTE deposition of DHB and DAN matrices onto mouse brain tissue. Representative ESEM images of the DHB surface layer at the gray and white matter parts of the brain cerebellum are depicted in Figure 1c, while Figure 1f showcases the morphology of DAN crystals on the surface layer of the coated cerebellum gray and white matter. These images reveal a homogeneous distribution of DHB and DAN crystals across the matrix surface layer.

Simultaneously, we explored the homogeneity of the matrix layer deposited by LTE onto silicon slides. Nine thickness measurements were acquired from different areas of the silicon slide coated with 240 µg/cm^2^ of DHB matrix (Supplementary Figure S3b), yielding an average thickness of 1609.22 nm (Supplementary Table S1). Notably, the standard deviation across the slide was as low as 2.54%. These results underscore the capability of LTE deposition to produce a very uniform matrix layer with a homogeneous distribution of matrix crystals over the surface layer.

Apart from the homogeneous distribution of matrix crystals over the surface layer, the size of these crystals also significantly impacts MALDI-MSI performance, as it can influence ionization efficiency. Larger crystals require higher laser power for ionization compared to smaller crystals^26^. Utilising ESEM, we calculated the size of DAN and DHB crystals deposited by the LTE method, revealing small DAN crystals with an average size of 0.34 µm and a standard deviation of 18.17% (Supplementary Figure S6a), while DHB matrix produced crystals with an average of 0.54 µm and a standard deviation of 23.36% (Supplementary Figure S6b). In the context of current advancements towards single-cell imaging^27,28^ with pixel sizes as small as 1-2µm^2^, LTE technique represents a significant advancement in matrix deposition for MALDI-MSI applications.

### 3.3. Influence of LTE and deposition protocol on matrix purity for MALDI MSI

Next, we investigated the efficacy of LTE combined with controlled source shutters during matrix evaporation to prevent the deposition of volatile impurities into tissue samples, thereby enabling the utilization of less pure matrices for MALDI-MSI studies. To this end, we deposited two types of DHB and DAN matrices differing in purity (non-pure <99% versus pure >99%) on empty glass slides using two distinct deposition protocols: controlled shutters (C_Sh) versus open shutters (O_Sh). The controlled shutters protocol, as described in Supplementary Methods, entails maintaining the shutters closed until the evaporation point of the matrix is reached, whereas the second protocol involves opening both the source and substrate shutters from the onset of deposition, allowing all evaporated particles from the organic matrix material to deposit on the substrate.

After MALDI-MSI analysis, we employed principal components analysis (PCA) to discern how the purity of the matrix and the deposition protocol influence the variance in the data (Figure 2), with each point representing a pixel from either sample. Approximately 2000 pixels were acquired per sample, and 1000 random pixels were selected for data analysis. When the shutters were left open during the deposition process (Figure 2a), we observed that the second principal component (PC2) effectively separated the two matrices for both DHB and DAN, with the primary variance in the dataset (approximately 40%) attributed to the purity of the matrix. In this scenario, the initial heating of the evaporator source containing the matrix to 55°C, gradually increasing to 80°C, may lead to the evaporation of volatile impurities with temperatures different from 80°C, consequently depositing impurities onto the substrate sample.

**Figure 2:**
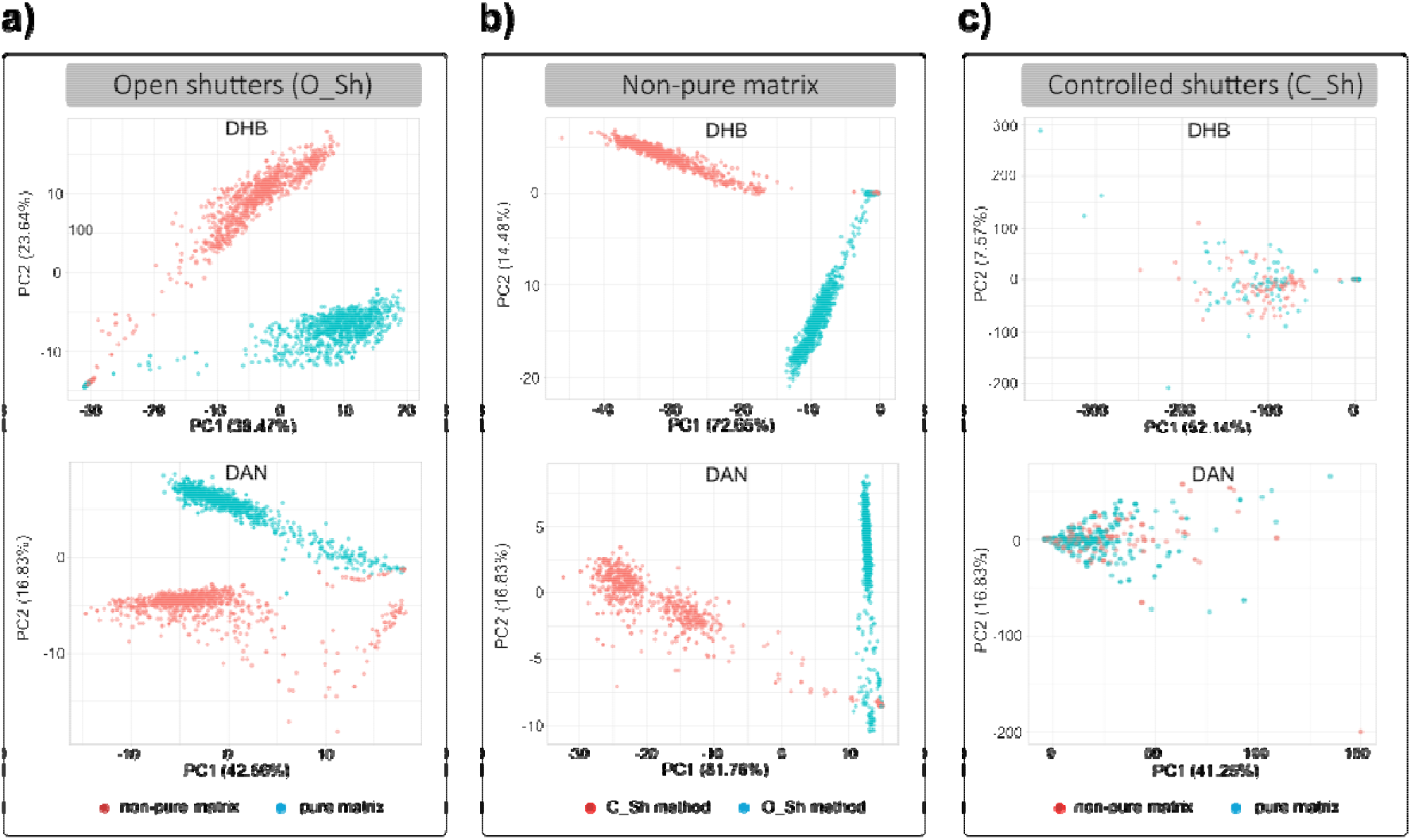
Deposition of pure and non-pure DHB (top) and DAN (bottom) matrices, using the open shutters (O_Sh) or controlled shutters (C_Sh) method, and related data from MALDI-MSI of only matrix. a) PCA plots of principal components 1 vs. 2 acquired using Open shutters method, showing separation between non-pure (red) and pure (blue) matrices. b) PCA plots of principal components 1 vs. 2 acquired using non-pure matrices with controlled shutters (red) vs. open shutters (blue). c). PCA plots of principal components 1 vs. 2 acquired using the controlled shutters method, illustrating overlapping between non-pure (red) and pure (blue) samples.

Subsequently, we examined the performance of non-pure DHB and non-pure DAN matrices using both the controlled shutter and open shutter approaches (Figure 2b), which exhibited separation by PC1 (explaining over 70% of the variance). As the same matrix purity was initially used for evaporation, the variance between the two groups stemmed from the deposition protocol.

Finally, we deposited pure and non-pure DHB and DAN matrices using our developed protocol with controlled shutters (Figure 2c). Notably, an overlap of the two samples (non-pure and pure matrices) was observed, indicating no differences between the two measured matrix sections. Unlike the open shutters method, opening the substrate shutters for matrix deposition when chamber temperatures reached 80°C effectively blocked other volatile impurities at temperatures below 80°C, preventing the desorption of impurities with temperatures higher than 80°C.

In summary, utilizing shutters to control when the matrix deposits on the sample allows the use of non-pure DHB and DAN matrices (approximately ten times cheaper than pure matrices) with comparable results.

Subsequently, we assessed the impact of matrix purity using our deposition protocol with controlled shutters on MALDI-MSI analysis of biological tissues. To this end, we compared the results of MALDI imaging of lipids from mouse brain sections when utilizing pure and non-pure DHB or DAN matrices (Supplementary Figure S7). This experiment was replicated three times (Supplementary Figure S8). Pooling all the results, we observed that the use of less pure matrices did not compromise lipid analytes ionization. Moreover, it preserved the high-resolution quality of the ion images when compared to those obtained from the pure matrices. This finding underscores the robustness and reliability of our deposition protocol with shutters in maintaining the integrity and effectiveness of MALDI-MSI analysis, irrespective of matrix purity.

### 3.4. Optimization and stability of LTE-deposited matrix layers for MALDI-MSI of brain tissue lipids

Initially, we optimized the matrix layer thickness for MALDI-MSI of lipids from mouse brain tissue. Using the calibration curves, we identified the thicknesses of 1.36 µm for DHB and 1.26 µm for DAN as optimal layers, providing stronger signals for intact lipids while minimizing matrix self-ionization (Supplementary Figures S9 and Fig S10).

Next, we investigated the stability of tissue samples coated with LTE-deposited matrix when stored at -80ºC. Consecutive mouse brain sections coated with DHB and DAN matrices were stored for 1 hour, 24 hours, and 14 days prior to MALDI-MSI analysis. Comparative analysis of the total ion counts (TIC) in each pixel from cerebellum brain sections at different time points is presented in Figure 3a and 3d. For both DHB and DAN matrices, stable analyte ionization was observed over the two-week storage period, as evidenced by comparable TIC and consistent average spectra. ESEM images of DHB and DAN matrices showed consistent small crystal distribution across storage periods of 1 hour, 24 hours, and 2 weeks, with stable average sizes (DHB: 0.54 µm; DAN: minor increase from 0.34 µm to 0.41 µm after two weeks) (Figure 3b and 3e) (Supplementary Figure S11). This stability was confirmed by MALDI-MSI results, which demonstrated identical lipid ion distribution patterns and comparable signal intensities across all storage durations (Supplementary Figures S12 and S13).

**Figure 3:**
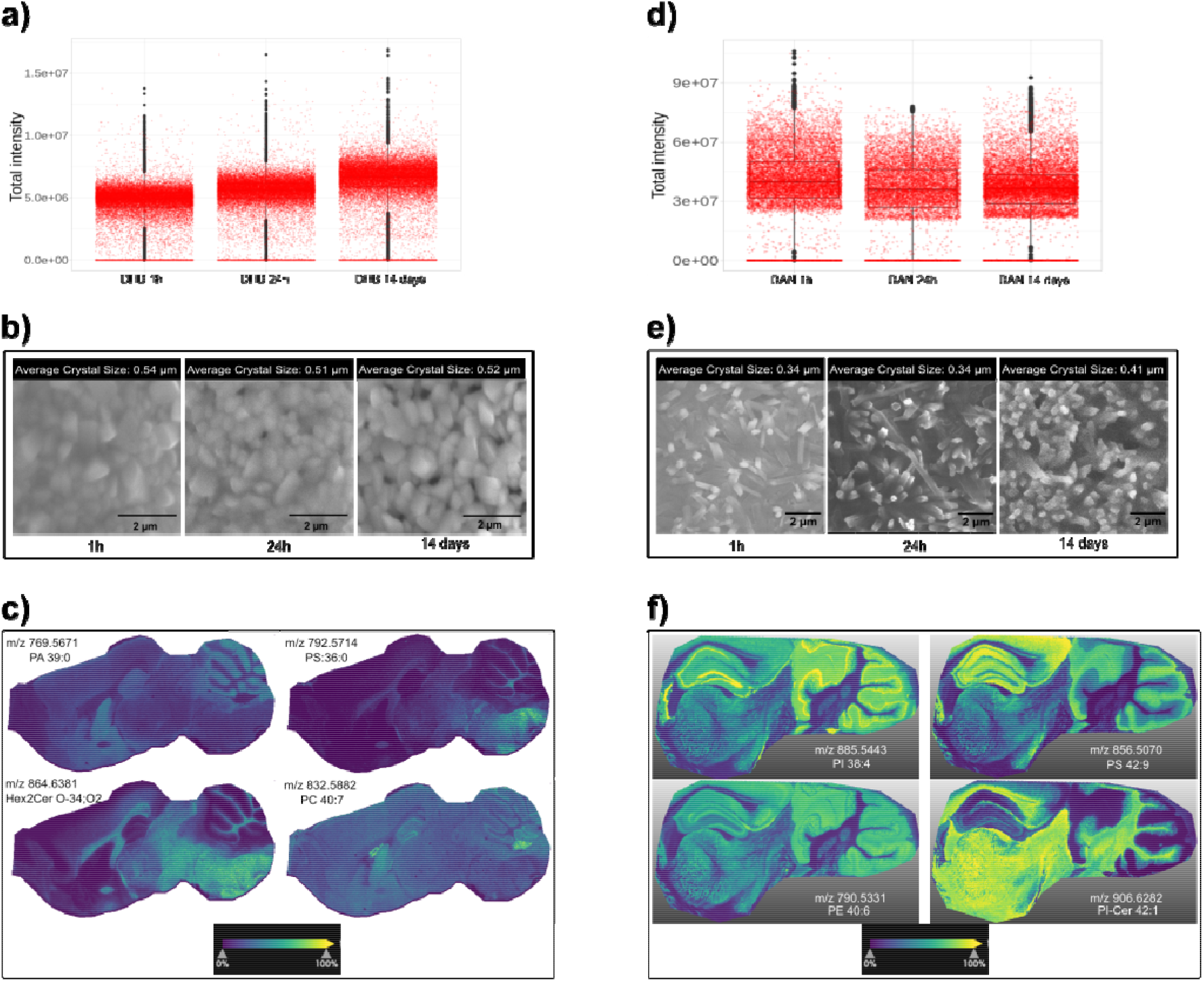
a) Boxplot showing the total ion count (TIC) intensity from mouse brain coated with DHB after storage at -80ºC for 1 hour, 24 hours, and 14 days. b) ESEM images of the DHB surface layer after storage durations of 1 hour, 24 hours, and 14 days; scale bar: 2 µm. c) MSI images of four intact lipids acquired at 20 µm lateral resolution from mouse brain coated with DHB matrix, using MALDI-Orbitrap mass spectrometer: PA 39:0 (*m/z* 769.5671, [M+Na]+), Hex2Cer 34:0;O2 (*m/z* 864.6381, [M+H]+), PS 36:0 (*m/z* 792.5714, [M+H]+), PC 40:7 (*m/z* 832.5885, [M+H]+). d) Boxplot representing the TIC intensity from mouse brain coated with DAN after storage at -80ºC for 1 hour, 24 hours, and 14 days. e) ESEM images of the DAN surface layer after storage durations of 1 hour, 24 hours, and 14 days; scale bar: 2 µm. f) MSI images of four intact lipids acquired at 10 µm lateral resolution from mouse brain coated with DAN matrix, using timsTOFfleX mass spectrometer: PI 38:4 (m/z 885.5443, [M-H]-), PE 40:6 (*m/z* 790.5331, [M-H]-), PS 42:9 (*m/z* 856.5070, [M-H]-), PI-Cer 42:1 (*m/z* 906.6282, [M-H]-).

To reinforce these findings, we performed high-resolution MALDI-MSI of mouse brain tissue using DHB and DAN matrices after storing the tissue at -80ºC for 15 and 23 days, respectively. Analysis with MALDI-Orbitrap and timsTOF flex equipment revealed exceptional image quality at resolutions of 20 µm and 10 µm, respectively, showcasing distinct lipid distribution patterns in various brain regions (Figure 3c and 3f). These results affirm that storing tissue coated with LTE-deposited matrices at -80ºC preserves ionization efficiency and image quality, even at high spatial resolutions, indicating the robustness of LTE-deposited matrices over time.

The thermal evaporator nanoPVD-T15A’s substrate holder accommodates three microscopic slides, enabling simultaneous matrix deposition onto three tissue slides in one run. Importantly, our investigation found that tissue slide positioning on the substrate area did not affect MALDI-MSI results (Supplementary Figure S14). This stability of coated tissue samples at -80ºC not only streamlines the experimental workflow by reducing the number of experimental depositions from three to one but also facilitates collaborative research by centralizing sample preparation and enabling the shipment of coated samples to other laboratories for MALDI-MSI acquisition if needed.

### 3.5. Comparative analysis of LTE and spray-coating methods in matrix deposition and MSI image quality for MALDI-MSI of mouse brain tissue

The optimized LTE method was compared directly to the spray method through MALDI-MSI using timsTOF-flex on serial sections of mouse brain for both DAN and DHB matrices. While matrix application with LTE was performed one month before MSI acquisition, the spray-coating samples underwent MSI analysis immediately after matrix application. Figure 4a and 4b illustrate the comparison of representative average spectra and TIC plots for DAN matrix deposition using LTE and spray methods, respectively. In the spray-coating data, signal intensities of ions detected in the lipid mass range (*m/z* 600 to 1000 Da) were lower compared to those obtained from LTE deposition. The TIC, representing the sum of ions detected in each pixel, showed an improvement with the LTE method over spray coating. After lipid annotation, the LTE samples revealed 176 annotated lipids, while the spray-coating method yielded 102. In Figure 4c, the lipid species annotated from mouse brain sections coated with DAN matrix are compared. Both deposition techniques exhibited high sensitivity in detecting glycerophosphoethanolamine (PE) lipid class, with 43 and 36 PE lipids annotated from LTE and spray-coating samples, respectively. Notably, the LTE method showed greater numbers of all other detected lipid classes compared to spray-coating, particularly lysophosphatidylcholine (LP), sphingolipids (SP), and glycerophosphate (PA) species.

**Figure 4:**
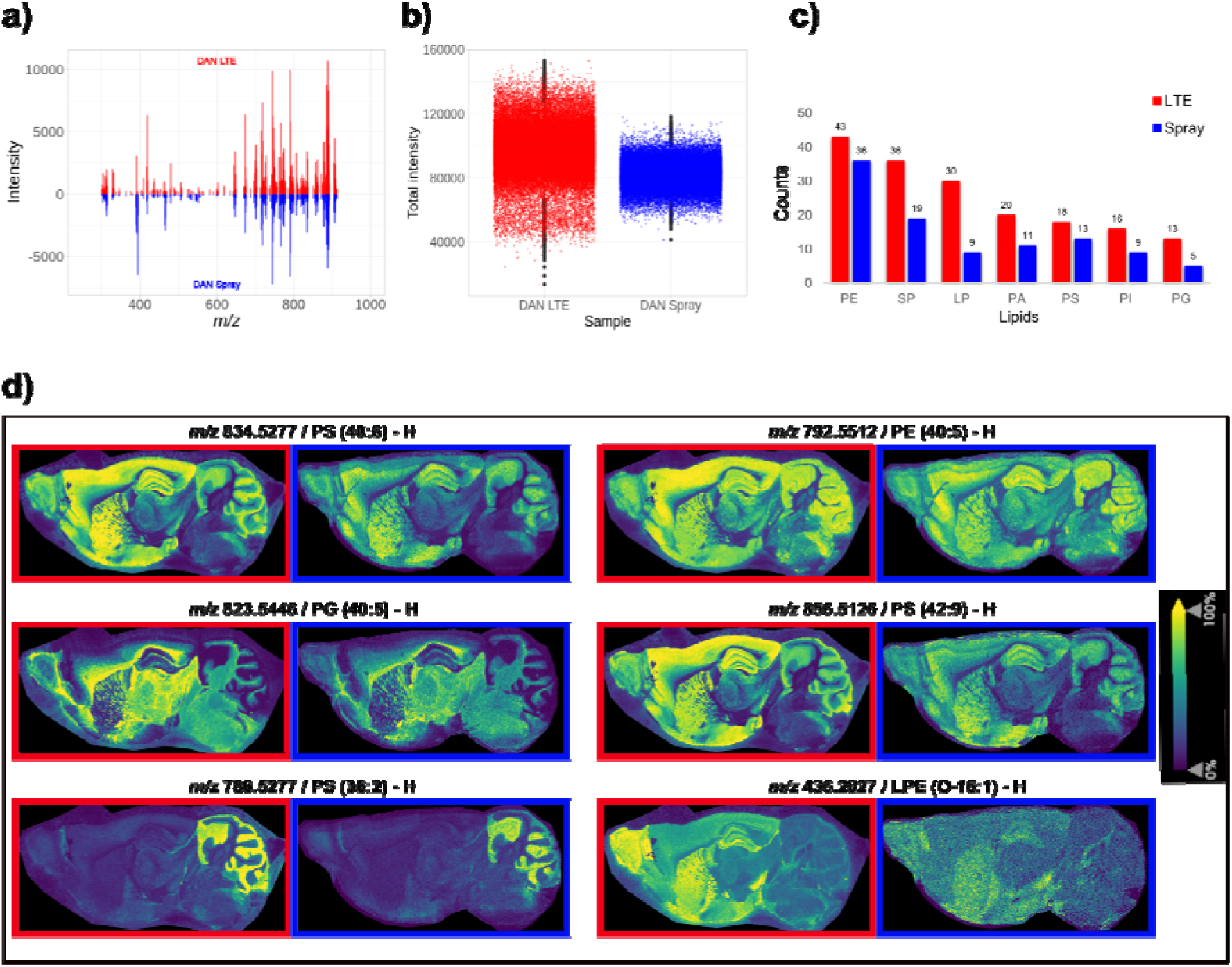
MALDI-MSI analysis of mouse brain tissue coated with DAN matrix using LTE deposition (red) and spray deposition (blue). a) Average spectra comparison. b) Boxplots representing the intensity of the total ion count (TIC) for each deposition method. c) Bar plot illustrating the number of annotated lipid species detected using LTE (red), spray method (blue). d) MSI images acquired at 30 µm lateral resolution using timsTOFfleX mass spectrometer, depicting six intact lipid ions: PS 40:6 (*m/z* 834.5277), PE 40:5 (*m/z* 792.5512), PG 40:0 (*m/z* 823.5446), PS 42:9 (*m/z* 856.5126), PS 36:2 (*m/z* 786.5277), and LPE O-16:1 (*m/z* 436.2527). All lipid ions were detected in negative mode ([M-H]^-^). Images obtained using LTE and spray techniques are delineated in red and blue, respectively.

Furthermore, we evaluated the quality of MSI images for lipid ions obtained from both deposition methods. Using DAN deposited via LTE and spray-coating, we presented six ion images of polar lipids from the sagittal mouse brain at 30 µm lateral resolution (Figure 4d). The MSI images from the LTE sample displayed clearer and sharper features compared to those from the spray-coating method. Notably, LTE deposition exhibited less lipid diffusion, evident in the pattern distribution of lipids such as PS 40:6 (*m/z* 834.5277), PE 40:5 (*m/z* 792.5512), and PS 42:9 (*m/z* 856.5126). In contrast, spray-coating resulted in lipid diffusion from the cerebellum and cerebral cortex towards the brain section periphery. Conversely, LTE delineated brain regions more clearly, including the molecular layer and granular layer of the cerebellum, emphasized by the distribution of lipid PS 42:9 (*m/z* 856.5126). This distinction was challenging with spray-coating due to analyte diffusion and delocalization. Furthermore, the distribution of lipid PS 36:2 (*m/z* 786.5277) highlighted the cerebellum’s structure, showing a homogeneous distribution with LTE deposition, unlike spray-coating, which exhibited varying abundances within the layer. MSI images obtained using spray-coating were blurred for low molecular weight lipids like LPE (*m/z* 436.2527), whereas LTE showed improved clarity and better-defined brain regions, such as the hippocampus.

To objectively quantify the quality of MSI images, we evaluated two aspects: the dispersion of intensity within LTE and spray-coating samples, and the accuracy of defining histological mouse brain regions using image segmentation.

Focusing on a comparable cerebellum region in both images, we represented pixel intensities using density plots for a selected group of ions (Supplementary Figure S15). Specifically, considering lipid PS 36:2 (*m/z* 786.5277), which highlighted the cerebellum region (Figure 4d), LTE deposition exhibited improved uniformity of signal intensity within the selected pixels (sd=0.0544) compared to the spray-coating method (sd=0.1095) (Supplementary Figure S15c).

Additionally, we compared the reference mouse brain atlas (Figure 5a) with the clustering outcomes of two MSI images obtained using LTE (Figure 5b) and spray-coating (Figure 5c) deposition of DAN matrix. For both methods, employing 6 clusters proved sufficient to delineate various anatomical regions of the mouse brain, including the cerebellum, hippocampus, isocortex, and the main olfactory bulb (MOB). Notably, images zoomed in using the LTE method exhibited finer structural details of the MOB (Figure 5b-1) and the hippocampus (Figure 5b-2), aligning better with the reference atlas compared to those obtained with the spray-coating method (Figure 5c-1 and 5c-2). Furthermore, segmentation of the sagittal brain coated with LTE (Figure 5b-3) showed excellent correspondence with the reference atlas, accurately delineating specific parts of the cerebellum (gray and white matter). Particularly, the molecular layer of the gray matter was distinctly defined with a single cluster (depicted in purple in Figure 5b-3). In contrast, clustering of the spray-coating image divided the molecular layer region into two clusters, with an additional yellow-colored cluster (Figure 5c-3), inconsistent with the cerebellum’s anatomical structure. Moreover, these cluster groups appeared outside the cerebellum and the isocortex region (near the edges). Similar observations were made with clustering using k=5 for the MSI image (Supplementary Figure S16). In LTE deposition, the analyte composition of the molecular layer region remained consistent, resulting in a homogeneous cluster. Conversely, the presence of two cluster groups in spray-coating results could be attributed to analyte diffusion and delocalization caused by the spray-coating method.

**Figure 5:**
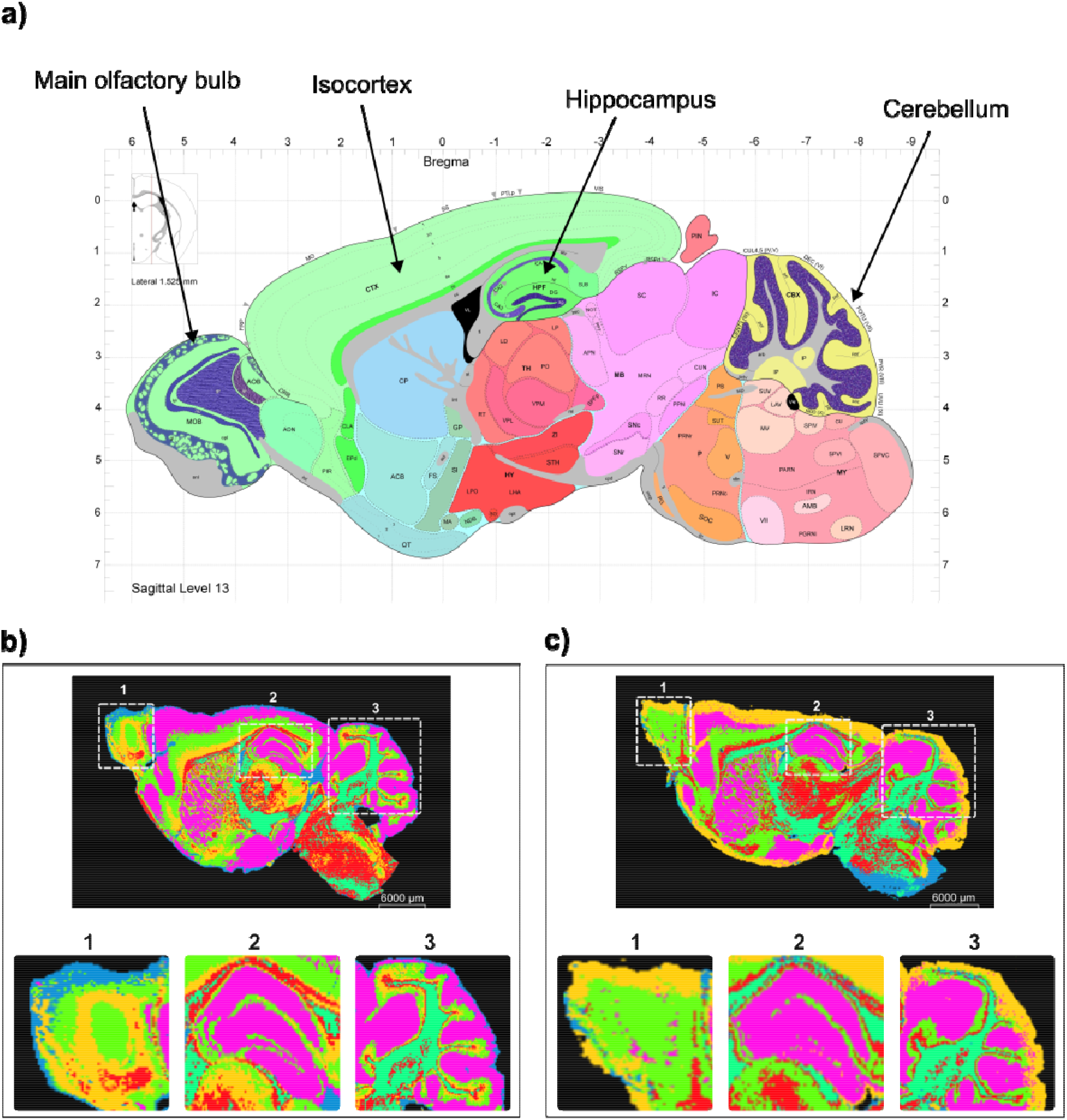
a) Anatomical reference atlas displaying sagittal mouse brain structures with a systematic, hierarchically organized taxonomy^29^. b) Segmentation outcomes of sagittal mouse brain tissue coated with DAN matrix using LTE deposition (top) with zoom-in images of the main olfactory bulb (b1), hippocampus (b2), and cerebellum (b3) regions. c) Segmentation results of sagittal mouse brain tissue coated with DAN matrix using spray-coating deposition (top) with zoom-in images of the main olfactory bulb (c1), hippocampus (c2), and cerebellum (c3) regions. Number of clusters=6; scale bar: 6000 µm.

## 4. Conclusions

We present a novel dry deposition method, low-temperature thermal evaporation (LTE), for the precise and reproducible application of thin organic matrices in MALDI-MSI. Our approach offers several advantages over existing methods. The high vacuum pressure achieved by the thermal evaporator nanoPVD-T15A enables matrix evaporation at a low temperature (80°C), significantly reducing sample contamination and biological degradation during deposition. Additionally, we demonstrate the ability to achieve a linear calibration of deposited matrix layers, allowing for precise control of thickness, essential for optimizing MALDI-MSI experiments. Notably, the LTE protocol serves as an additional purification step, blocking impurities with shutters during deposition, resulting in very pure matrix layers irrespective of matrix quality. Furthermore, LTE deposition yields homogeneous layers of small-sized matrix crystals (average size: 0.54 µm for DHB and 0.34 µm for DAN) on tissue surfaces. Importantly, the morphology and crystal size of DHB and DAN matrices remain stable when stored at -80ºC for at least 2 weeks, with no impact on ionization efficiency or MSI image quality. Current advancements are steering towards single-cell imaging with pixel sizes as small as 1-2µm^2^, driven by instrumental progress. Our LTE technique holds promise in enhancing MSI data quality owing to crystal sizes (< 1 µm), surpassing the pixel size. Comparative analysis with the spray-coating method demonstrates that LTE application of DAN matrix detects a larger number of lipids, exhibits higher signal intensities, reduces analyte diffusion, and improves MSI image quality. Overall, our LTE method represents a significant advancement in matrix deposition for MALDI-MSI applications, offering enhanced performance and reproducibility.

## Acknowledgements

TM acknowledges the financial support of the Universitat Rovira i Virgili through the predoctoral grant ref. PRE2019-089374. XC acknowledges the financial support of the Spanish Ministry of Economy and Competitivity through project RTI2018-096061-B-100. OY acknowledges the financial support of grant PID2022-136226OB-I00 funded by MICIU/AEI/10.13039/501100011033 and by ERDF/EU, and grant TED2021-132635B-I00 funded by MICIU/AEI/10.13039/501100011033 and by the European Union NextGenerationEU/PRTR. SAM acknowledges the financial support of DAAD Scholarship Funding programme/-ID: Research Grants - Doctoral Programmes in Germany, 2020/21 (57507871). CH acknowledges the funding by the German Federal Ministry of Education and Research (BMBF) as part of the MSCorSys SMART-CARE (grant 890 161L0212F) for the acquisition of the timsTOFflex, and the Innovation Partnership M2Aind, projects ADCtox-3D (13FH8E01IA) and Drugs4Future (13FH8I05IA).

