## Supporting Information for "Improving MALDI Mass Spectrometry Imaging Performance: Low-Temperature Thermal Evaporation for Controlled Matrix Deposition and Improved Image Quality"

**Supplementary Methods**

**Protocol for deposition of DHB and DAN on animal tissue cryosections by low temperature thermal evaporation (LTE) using the nanoPVD-T15A**:

1. Open the chamber and the sample shutter to place the sample slide facing down on the sample holder. Secure the sample using electrical tape. Up to three standard microscopy slides (75 x 25 mm) can fit in the sample holder.
2. Open the source shutter and fill the crucible with the chosen matrix powder, filling about ¾ of the crucible. Weigh the crucible with the matrix on a microbalance before placing it in the source. Place the crucible in the source and close the source cover properly, then close the source shutter.
3. Close the chamber and start the pumping process. Pump until the pressure is below $5x10^{-5}$ mbar. Wait for 3 minutes to ensure stable pressure after pumping. Pumping usually takes 2 minutes for DHB or DAN.
4. Set the initial temperature of the source to 55ºC and wait 10 minutes for temperature stabilization.
5. Increase the temperature slowly in 5ºC steps until reaching 70ºC. Ensure that power SP reaches 0 before increasing the temperature.
6. Increase the temperature from 70 to 76ºC with a 2ºC step, and from 76 to 80ºC with a 1ºC step. Open the source shutter when the temperature reaches 80ºC.
7. Monitor the quartz sensor and wait at least 5 min for rate stabilization. Then, open the sample shutter and start the rotation of the sample holder.
8. Monitor the desired thickness by the quartz sensor in Angstrom (Å). Once the desired thickness is reached, close the source and substrate shutters.
9. Set the temperature to 0ºC and vent the chamber. Venting the system usually takes 30 minutes for DHB or DAN and can only be done if the temperature is below 100ºC.
10. Once the chamber is vented, open the sample and source shutters, and carefully remove the sample and the crucible with the matrix.

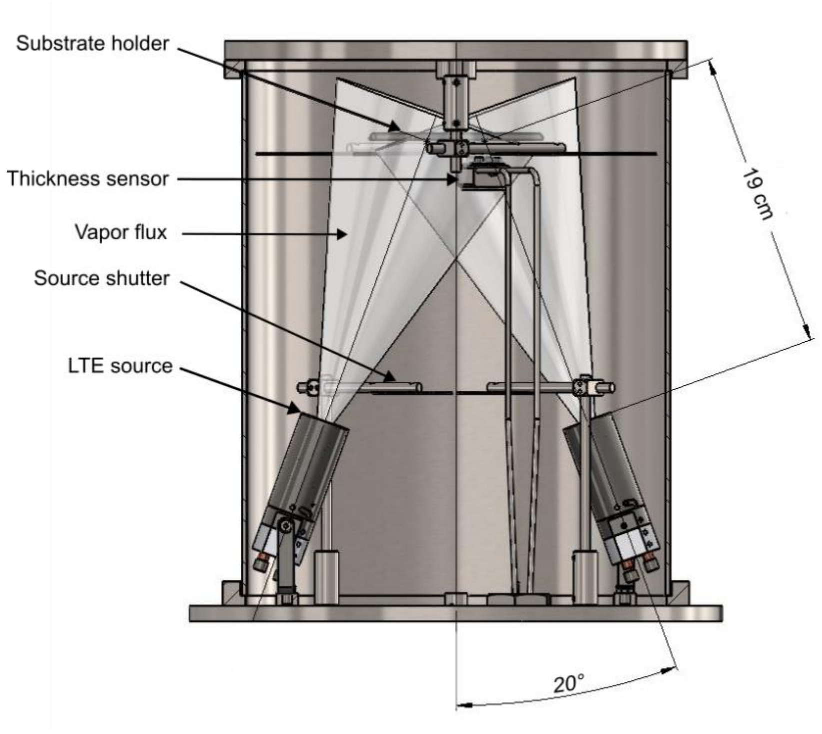


**Figure S1.** Schematic diagram of a nanoPVD-T15A process chamber with both LTE sources, source shutters, rotating substrate holder, and a thickness sensor. The distance between the source and the substrate is 19 cm, and the angular position of the sources is 20°.

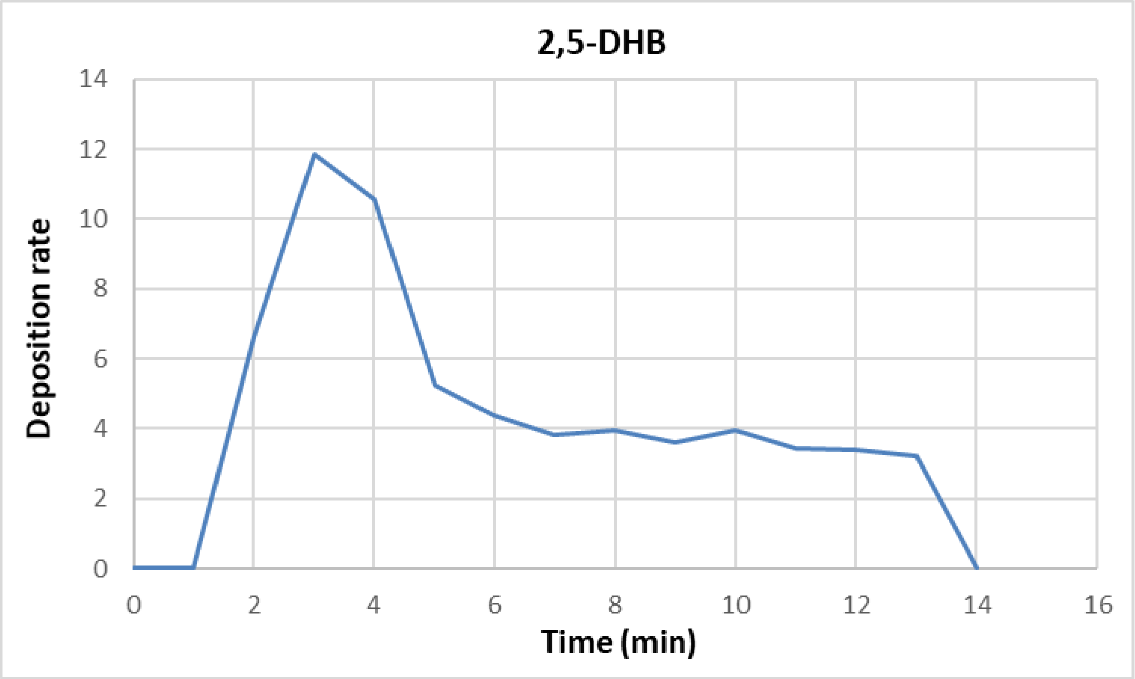


**Figure S2:** Graphical representation of the deposition rate, shown as the thickness of the deposited layer over time (Å/min), for the evaporated DHB organic matrix at 80°C, as measured by the thermal evaporator sensor. As the thermal source is heated to the optimal evaporation temperature, the deposition rate increases sharply, reaching a peak before slightly decreasing and stabilizing. This stable deposition rate is maintained throughout the deposition process.

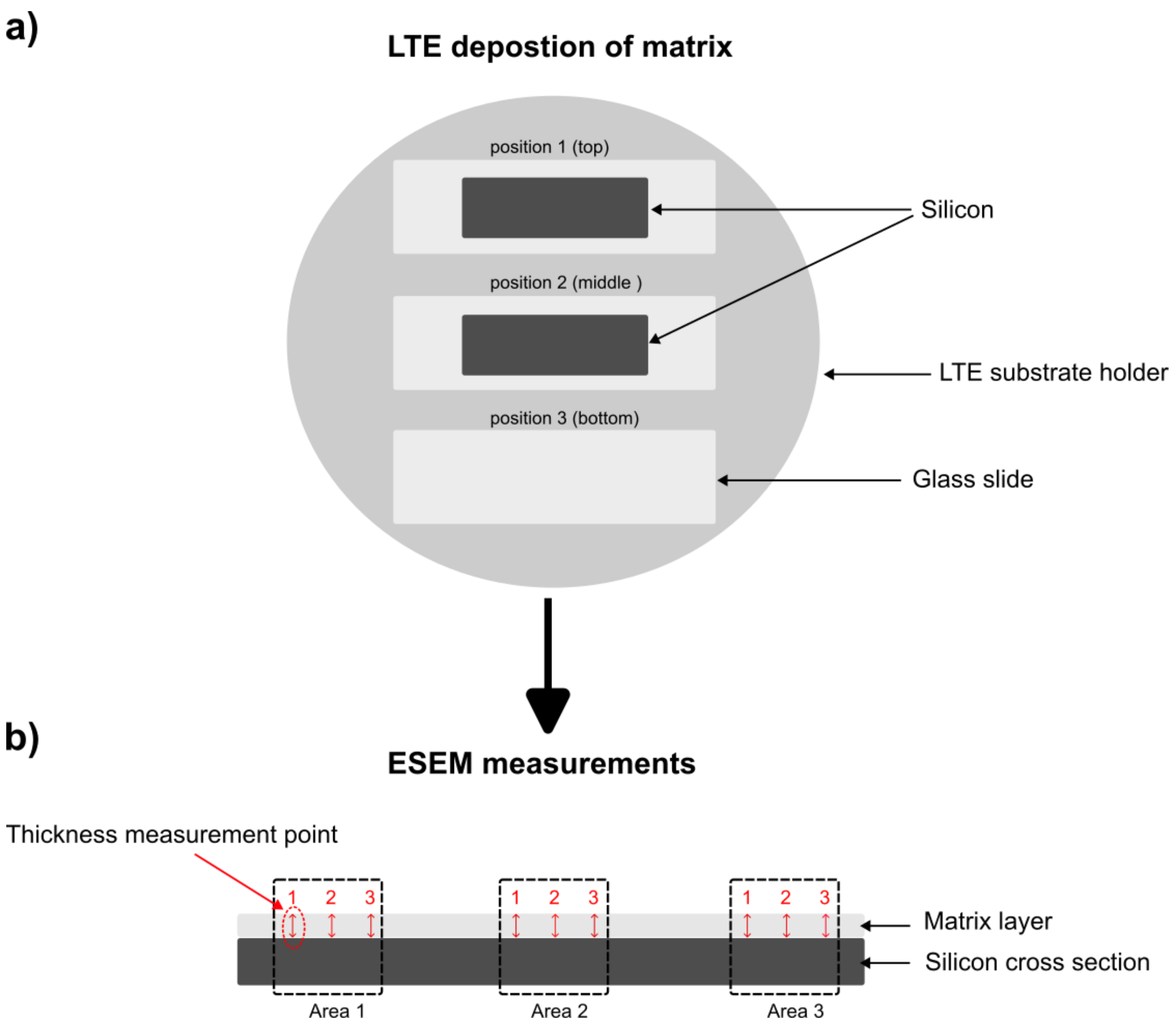


**Figure S3:** a) Schematic diagram illustrating the experimental setup for matrix deposition using LTE on silicon and glass slides. Two silicon slides were positioned at the top and middle of the substrate holder for subsequent ESEM measurements, while one glass slide was positioned at the bottom for measuring matrix coverage (µg/cm²).
b) Schematic diagram depicting the measurement of layer thickness using ESEM for each replicate.

**Table S1:** ESEM measurements were conducted on three distinct regions of a cross-sectioned silicon slide to assess the DHB matrix layer. This involved three measurements in each region, totaling nine measurements.

| Sample | Thickness monitored by the sensor (Å) | Thickness measured by ESEM (nm) | Average Thickness (nm) | Relative standard deviation (RSD%) |
|---|---|---|---|---|
| S_6 | 17500 | 1538 | 1609.22 | 2.54 |
| | | 1603 | | |
| | | 1581 | | |
| | | 1678 | | |
| | | 1673 | | |
| | | 1597 | | |
| | | 1597 | | |
| | | 1608 | | |
| | | 1608 | | |

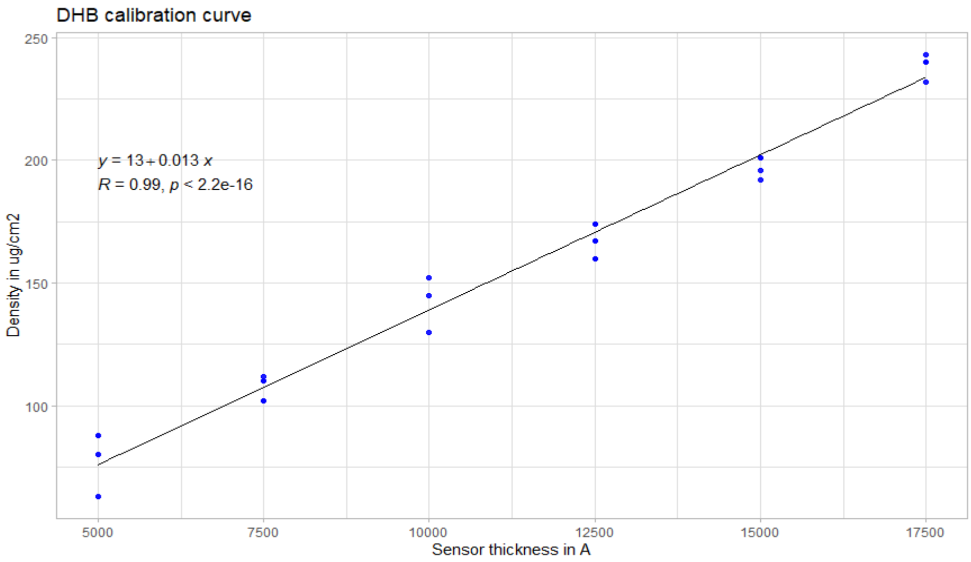


**Figure S4:** Calibration curve illustrating the deposition of DHB, correlating the sensor thickness reported in Å with the amount of matrix deposited on the slide area (µg/cm2).

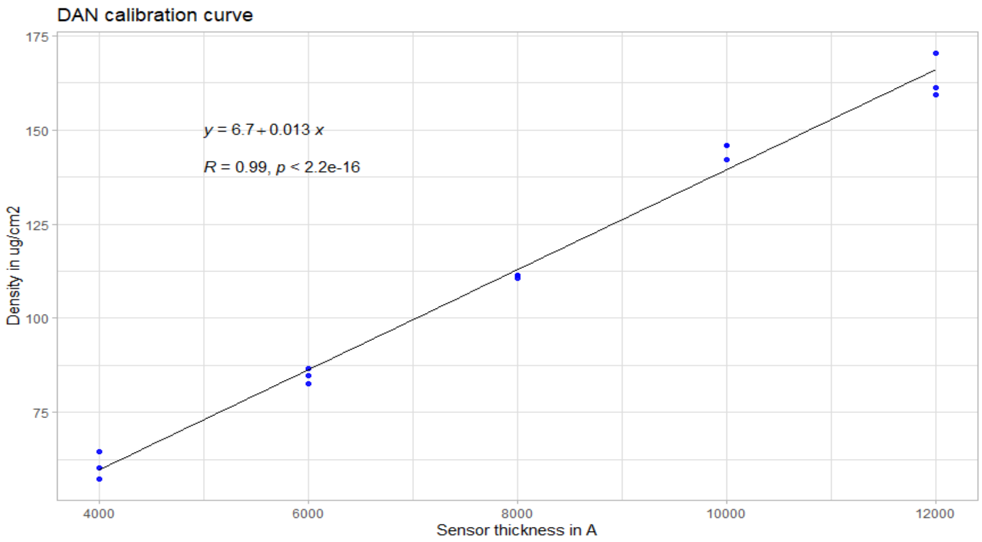


**Figure S5:** Calibration curve illustrating the deposition of DAN, correlating the sensor thickness reported in Å with the amount of matrix deposited on the slide area (µg/cm2).

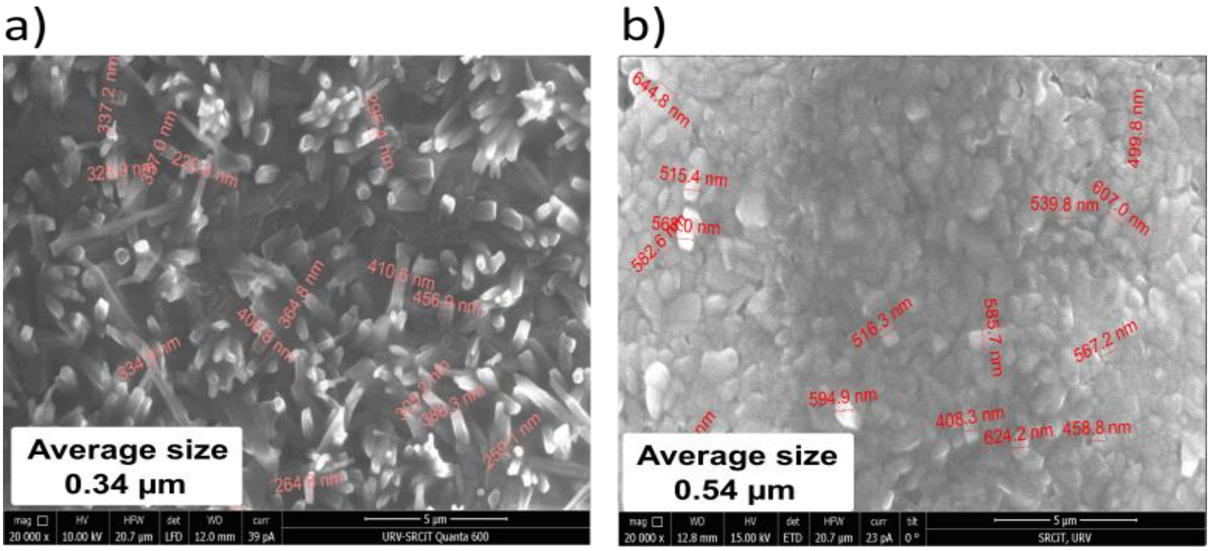


**Figure S6:** Environmental scanning electron microscopy (ESEM) images obtained at a magnification of 20,000 showing the surface of the matrix layer, along with crystal size measurements for both DAN (a) and DHB (b). Scale bar indicates 5 µm.

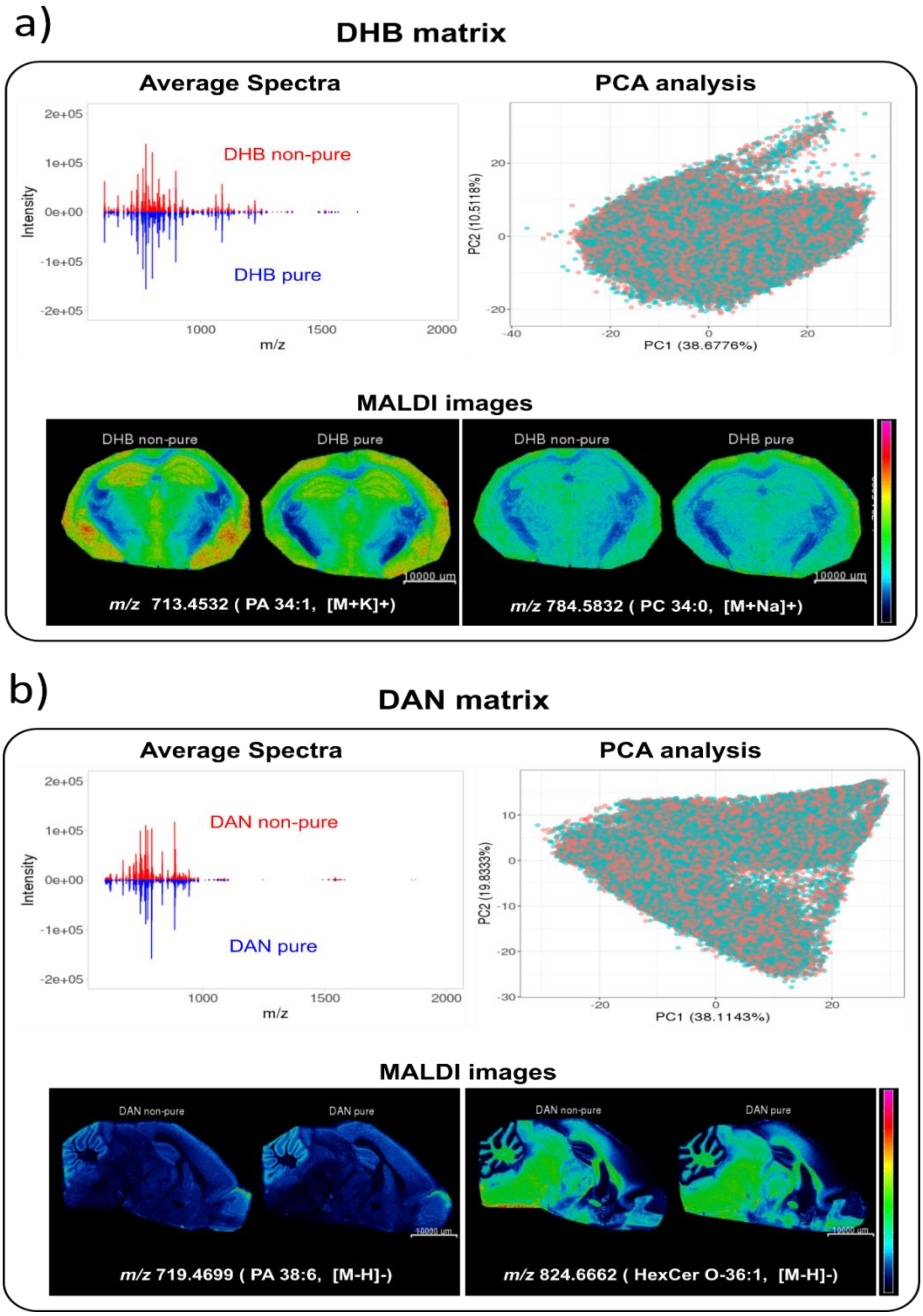


**Figure S7:** Comparison of non-pure and pure DHB and DAN matrix depositions and MALDI-MSI analysis. (a) MALDI-Orbitrap MSI results in positive mode using DHB: Average spectrum (top left) and Principal Component Analysis score plot (top right) of mouse brain samples coated with non-pure DHB (red) and pure DHB matrix (blue). Two examples of intact lipids detected by MSI: *m/z* 713.4532 (glycerophosphate PA 34:1, [M+K]+), and *m/z* 784.5832 (glycerophosphocholine PC 34:0, [M+Na]+). (b) MALDI-Orbitrap MSI results in negative mode using DAN: Average spectrum (top left) and Principal Component Analysis score plot (top right) of mouse brain samples coated with non-pure DAN (red) and pure DAN matrix (blue). Two examples of intact lipids detected by MSI: *m/z* 719.4699 (phosphatidylethanolamine, [M-H]-) and *m/z* 824.6662 (3-O-Sulfogalactosylceramide, [M-H]-).

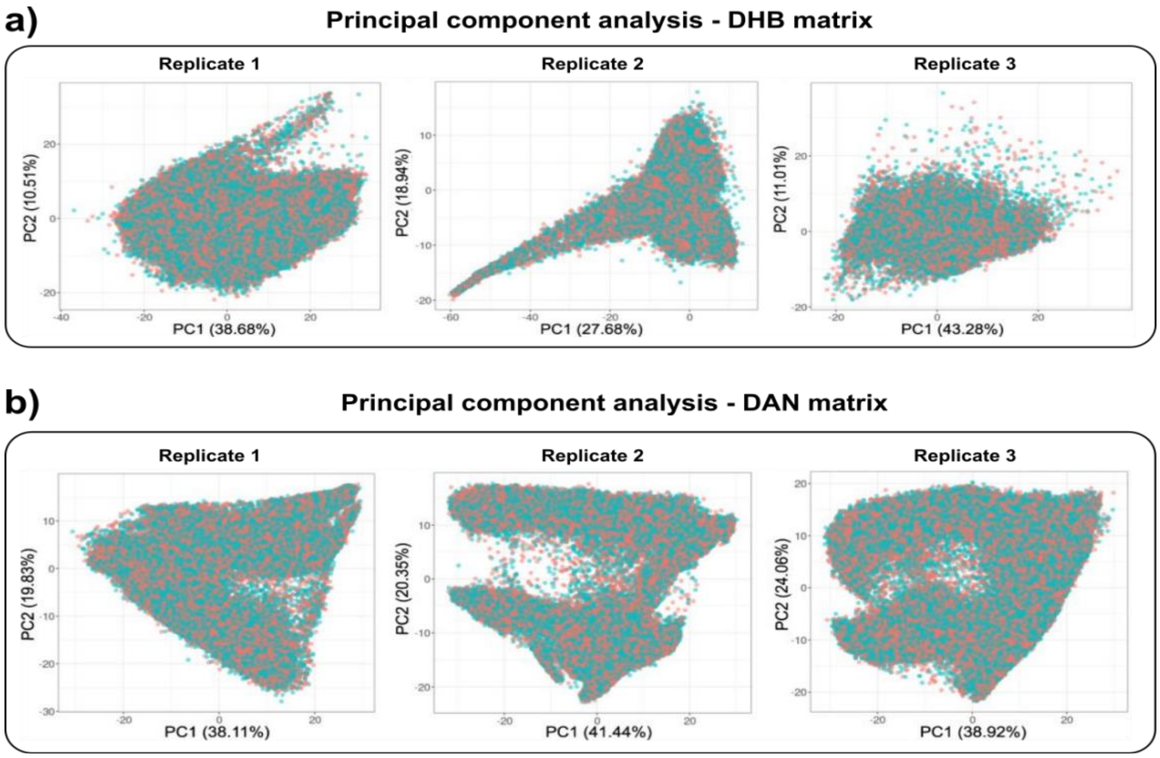


**Figure S8:** a) MALDI-Orbitrap MSI analysis of mouse brain coated with pure (>99%) and non-pure (98%) DHB matrix. PCA plots of principal components 1 vs. 2 of the DHB non-pure sample (red) and pure sample (blue) of three replicates. b) MALDI-MSI analysis of mouse brain coated with pure (>99%) and non-pure (97%) DAN matrix. PCA plots of principal components 1 vs. 2 of the DAN non-pure sample (red) and pure sample (blue) of three replicates.

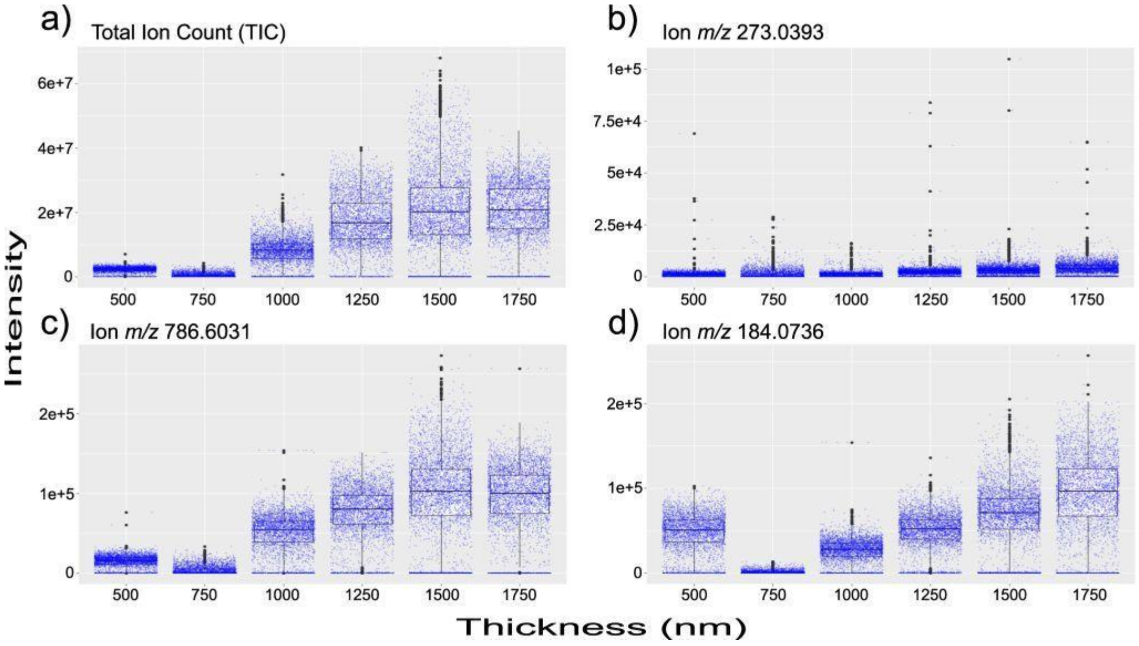


**Figure S9:** Boxplots depicting the intensity of signals detected on homogenized liver tissue coated with six layers of DHB and measured in positive mode. Panels (a), (b), (c), and (d) show the intensity of the following ions, respectively: (a) total ion count; (b) matrix-related ion at *m/z* 273.0399; (c) $[M+H]^+$ of the glycerophosphocholine C44H84NO8P at *m/z* 786.6031; and (d) $[M+H]^+$ of the phosphocholine head-group at *m/z* 184.0736.

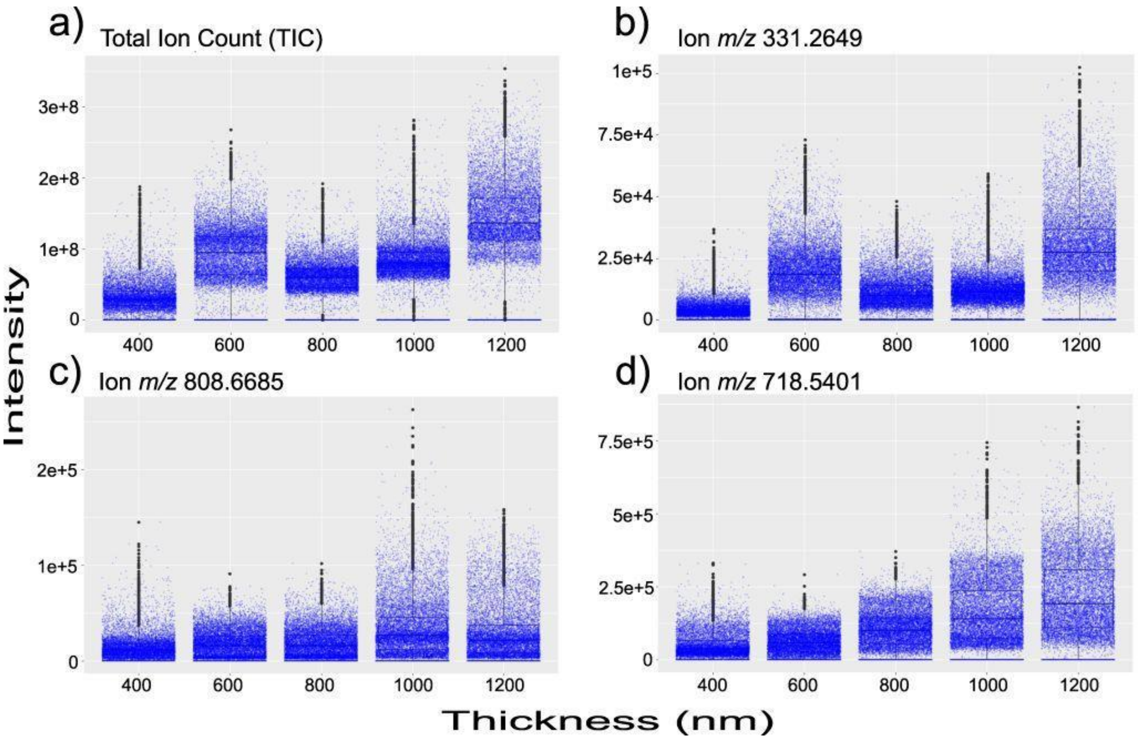


**Figure S10:** Boxplots illustrating the intensity of signals detected on brain tissue coated with five layers of DAN and measured in negative mode. Panels (a), (b), (c), and (d) show the intensity of the following ions, respectively: (a) total ion count; (b) matrix-related ion at *m/z* 331.2649 ($[M-H]^-$); (c) sphingolipid (C48H91NO8) at *m/z* 808.6685 ($[M-H]^-$); and (d) phospholipid (C39H78NO8P) at *m/z* 718.5401 ($[M-H]^-$).

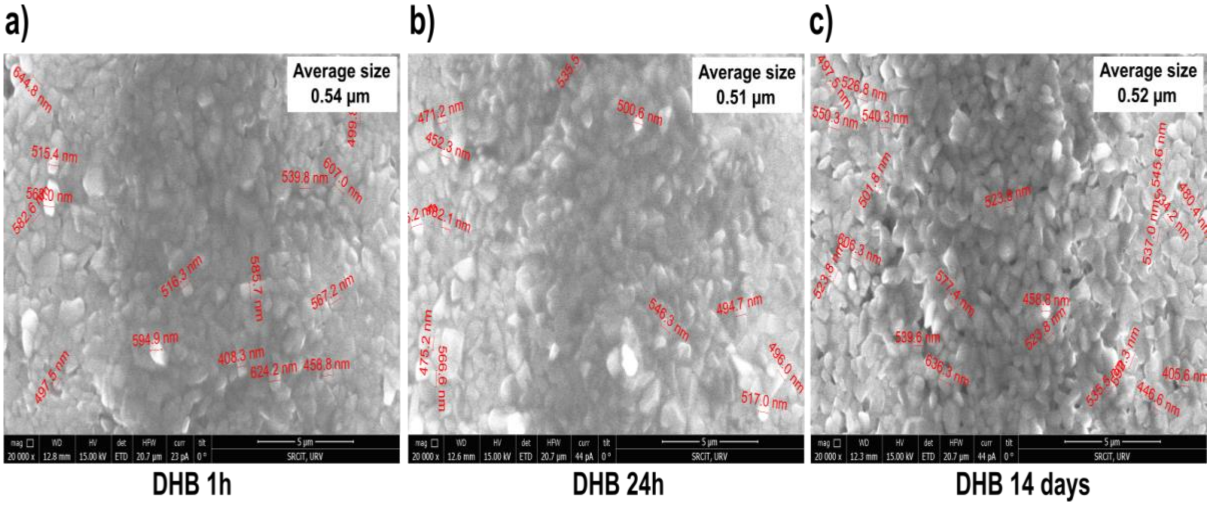


**Figure S11:** Environmental scanning electron microscopy (ESEM) images acquired at 20000 magnifications of the DHB matrix layer surface on top of the brain section, with crystal size measurements: (a) Coated tissue stored for 1 hour. (b) Coated tissue stored for 24 hours. (c) Coated tissue stored for 14 days. Scale bar, 5 µm.

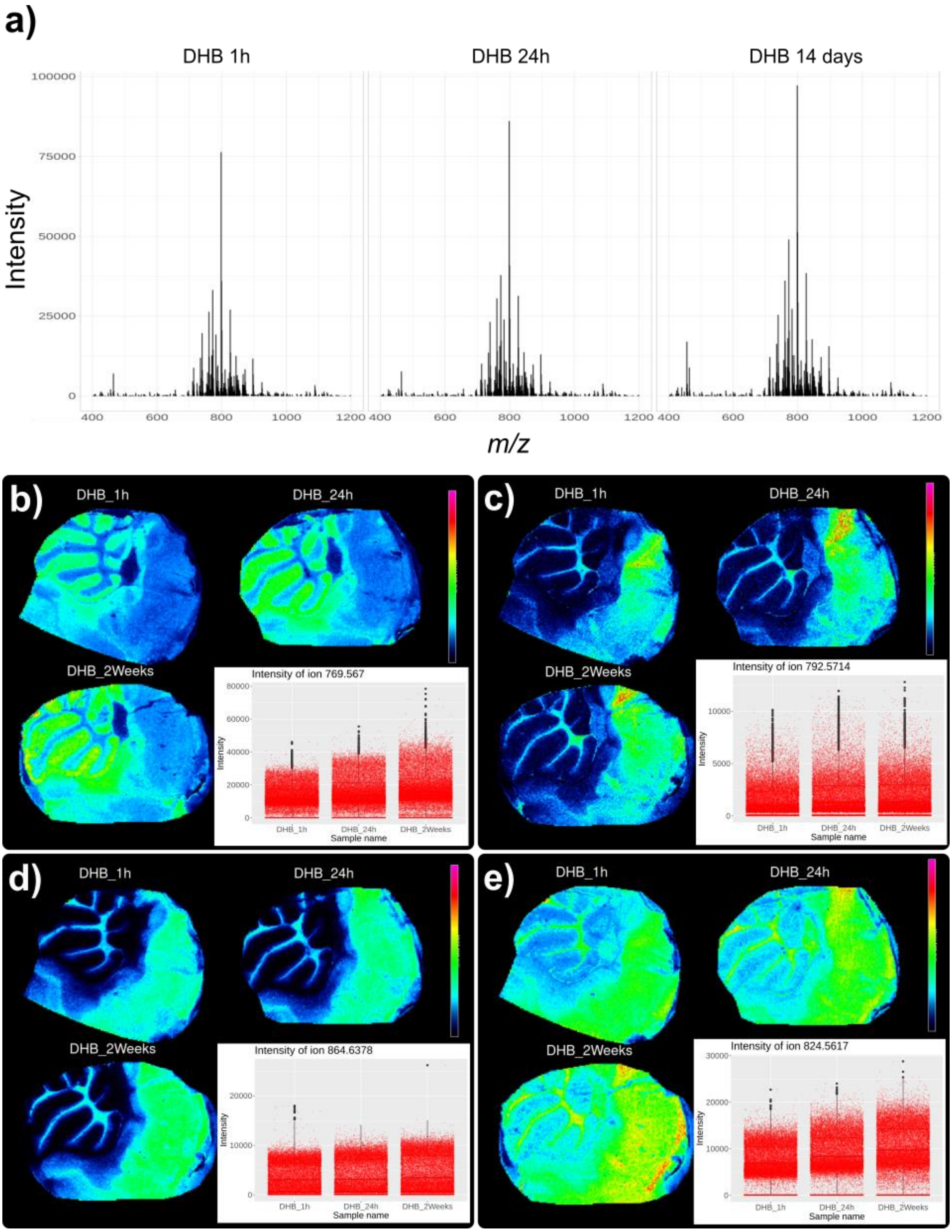


**Figure S12:** (a) Representative average spectra acquired from mouse brain tissue coated with DHB and stored at -80°C for varying durations: one hour (1h), twenty-four hours (24h), and two weeks (14 days). MALDI-Orbitrap MS images with corresponding intensity plots obtained from the three samples (1h, 24h, and 14 days) of ions are depicted. (b) Ion at *m/z* 769.5670 ([M+Na]+, PA 39:0). (c) Ion at *m/z* 792.5714 ([M+H]+, PS 36:0). (d) Ion at *m/z* 864.6378 ([M+H]+, Hex2Cer 34:0). (e) Ion at *m/z* 824.5617 ([M+K]+, PC 36:1).

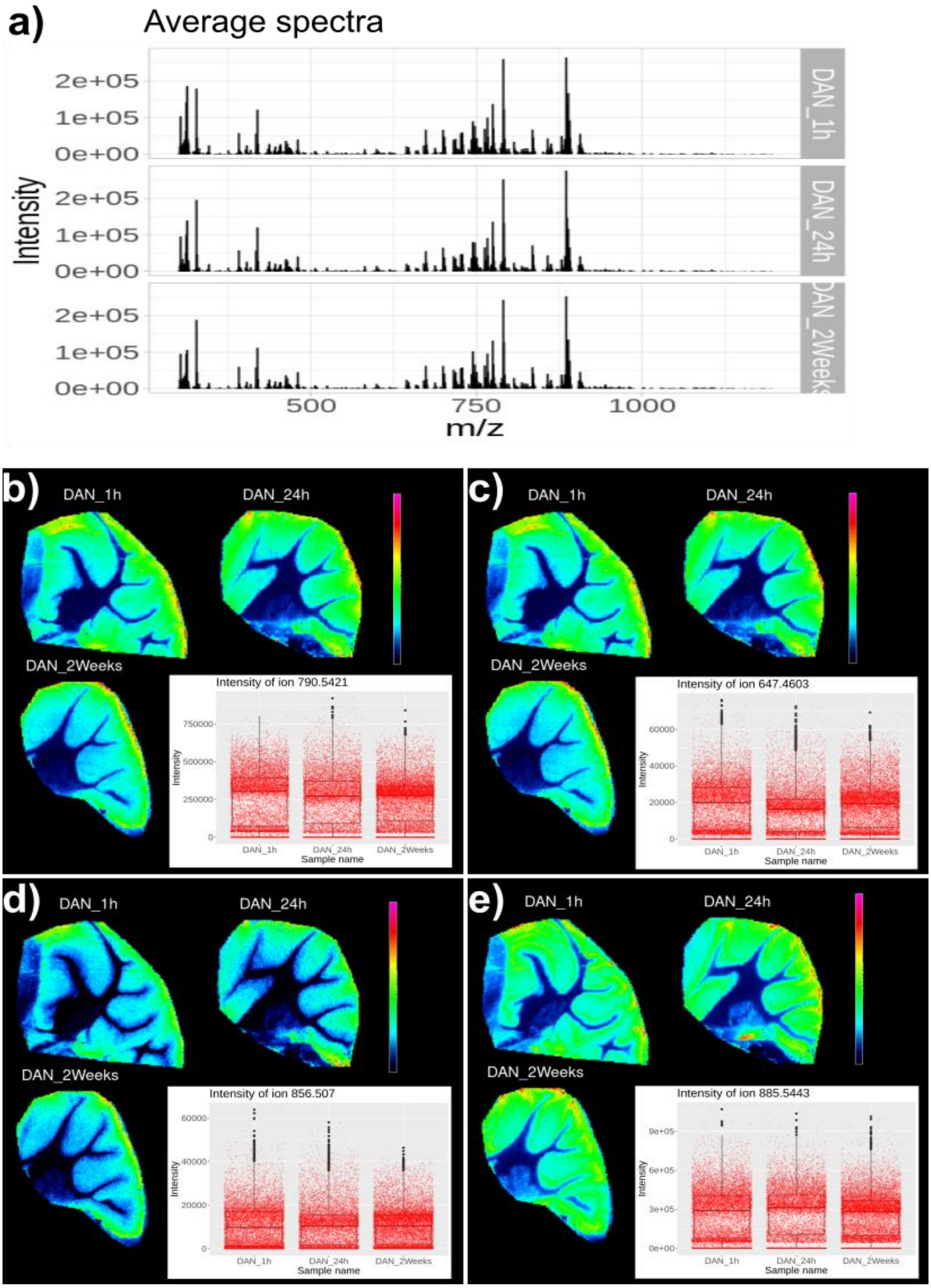


**Figure S13:** (a) Representative average spectra acquired from mouse brain tissue coated with DAN and stored at -80°C for varying durations: one hour (1h), twenty-four hours (24h), and two weeks (14 days). MALDI-Orbitrap MS images with corresponding intensity plots obtained from the three samples (1h, 24h, and 14 days) of ions are illustrated. (b) Ion at *m/z* 790.5421 ([M-H]-, PE 40:6). (c) Ion at *m/z* 647.4603 ([M-H]-, PA 32:0). (d) Ion at *m/z* 856.5070 ([M-H]-, PS 42:9). (e) Ion at *m/z* 885.5443 ([M-H]-, PG 41:5).

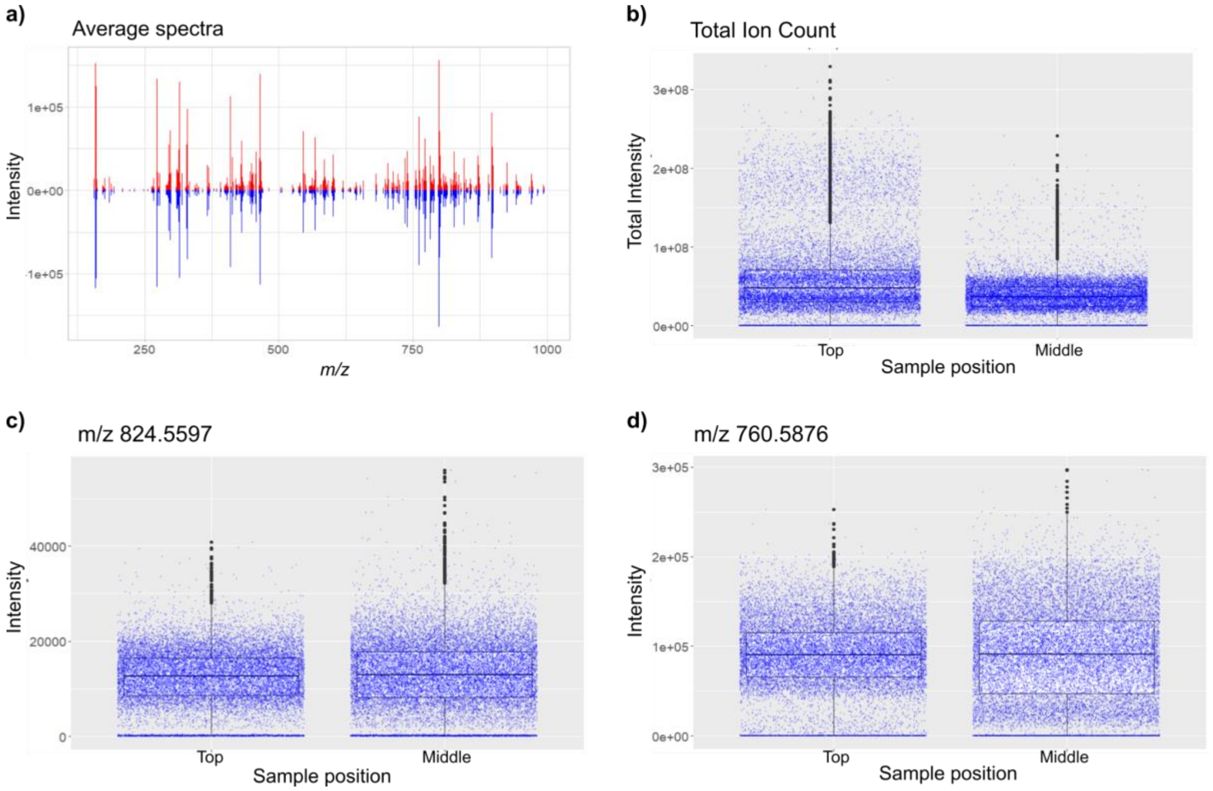


**Figure S14:** MALDI-MSI analysis in positive mode of two mouse cerebellum sections coated with DHB; one is placed on the top position of the slide holder, and the second on the middle. (a) Representative average spectra. (b) Boxplots representing the intensity of total ion count. (c) Boxplots representing the intensity of ion *m/z* 824.5597 ([M+K]+ of a Glycerophosphocholine (C44H84NO8P)). (d) Boxplots representing the intensity of ion *m/z* 760.5876 ([M+H]+ of a phospholipid (C42H82NO8P).

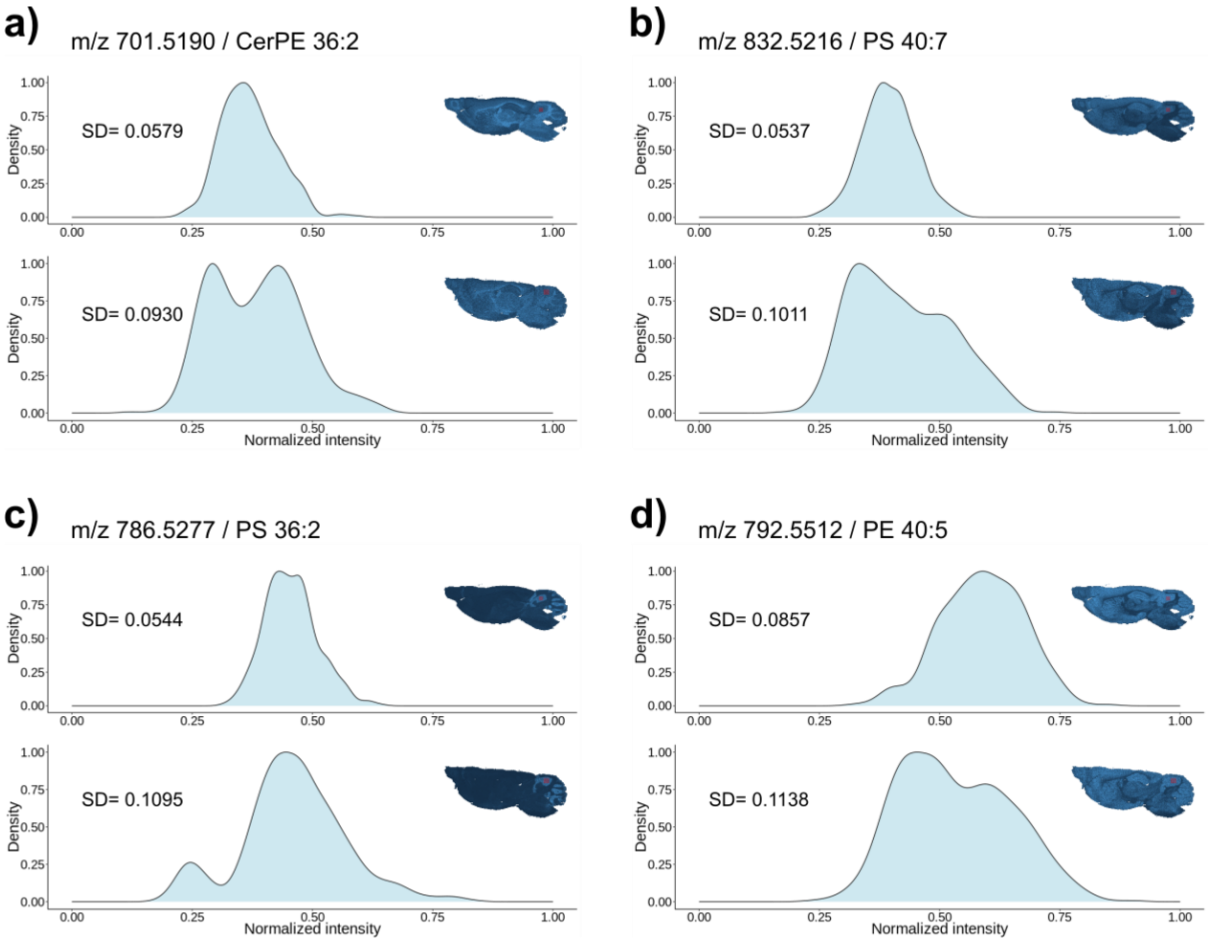


**Figure S15:** Representative histogram plots of the signal intensities (top: LTE, bottom: spray) of the selected mouse cerebellum region (rectangle area in red) with corresponding calculated standard deviation (SD): a) Ion at *m/z* 644.4964 ([M-H]-, CerP 36:1). b) Ion at *m/z* 647.4612 ([M-H]-, PA 32:0). c) Ion at *m/z* 786.5277 ([M-H]-, PS 36:2). d) Ion at *m/z* 792.5512 ([M-H]-, PE 40:5).

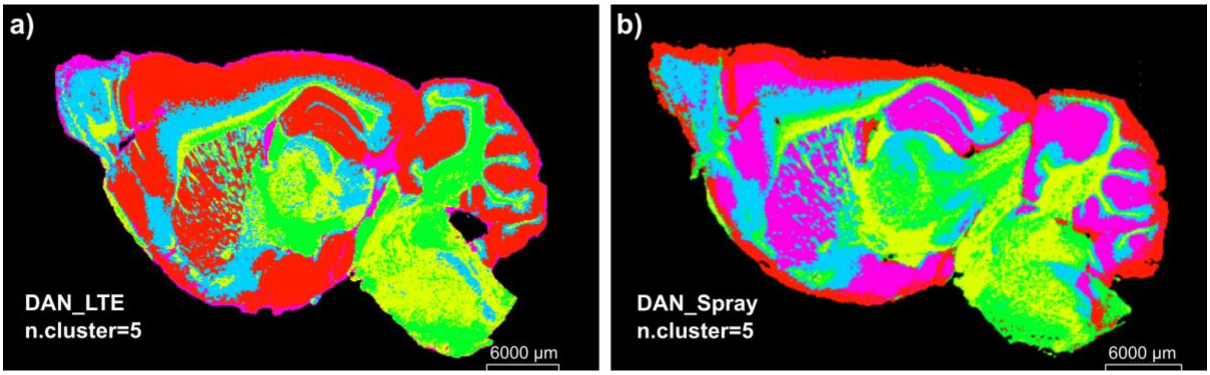


**Figure S16:** b) Segmentation results of the sagittal mouse brain tissue coated with DAN matrix using LTE deposition (top) with the zoom-in images of the main olfactory bulb (b1), hippocampus (b2), and cerebellum (b3) regions of the brain. c) Segmentation results of the sagittal mouse brain tissue coated with DAN matrix using LTE deposition (top) with the zoom-in images of the main olfactory bulb (b1), hippocampus (b2), and cerebellum (b3) regions of the brain; Number of clusters=6; scale bar: 6000 µm.
